# AptBacterialDB: A Comprehensive, Manually Curated Database of Antibacterial Aptamers

**DOI:** 10.64898/2026.07.01.735956

**Authors:** Nisha Bajiya, Ishika Gupta, Gajendra P. S. Raghava

## Abstract

In recent years, aptamers have transitioned from mere laboratory tools to highly potent molecular recognition agents capable of overcoming the strict limitations of conventional antibiotic therapies. We have developed AptBacterialDB, a manually curated, large, comprehensive database of experimentally validated antibacterial aptamers spanning 1996 to 2026. The database contains a total of 2131 aptamers targeting approx 75 different bacterial classes, and 124 aptamer targets with 95 entries found in UTexas databases, 97 in AptaDB, and 28 in Aptabase. It contains 1555 unique aptamer sequences, 189 unique modifications, 40 different selection approaches, and 44 different affinity methods. It integrates detailed annotations of about 20 fields, including sequence information, nucleic acid type, binding affinity, modifications, experimental and functional details. The secondary structure of the aptamers was predicted using ViennaRNA Package 2.0, demonstrating that they adopt mostly stable conformations, with a structured stem region. MySQL was implemented for database development, and a knowledge graph was integrated using ArcadeDB/openCypher for graphical visualization of aptamer-target-organisation relationships. Facilities such as different search modes, browsing, similarity search, REST API access, and entries linked to the existing database for a broader view of the aptamers have been provided. AptBacterialDB (https://webs.iiitd.edu.in/raghava/aptbacterialdb/) provides a user-friendly centralized platform to accelerate antibacterial aptamer research, therapeutic development, biosensor design, and computational modelling efforts.

**Author’s Biography:**

1. Nisha Bajiya is currently working as Ph.D. Student in Computational Biology from Department of Computational Biology, Indraprastha Institute of Information Technology, New Delhi, India.
2. Ishika Gupta is currently working as an M.Tech Student in Computational Biology from Department of Computational Biology, Indraprastha Institute of Information Technology, New Delhi, India.
3. Gajendra P. S. Raghava is currently working as a Professor in the Department of Computational Biology, Indraprastha Institute of Information Technology, New Delhi, India

## Introduction

Bacterial infections continue to pose a serious global health burden, further exacerbated by the rapid emergence of multidrug-resistant and extensively drug-resistant strains. Antimicrobial resistance (AMR) has emerged as one of the most severe global health threats of the twenty-first century. A report by a systematic study in 2021 estimated that nearly 4.71 million deaths were associated with bacterial AMR, with 1.14 million deaths directly attributable to resistant infections. The study also projects that by 2050, around 1.91 million deaths may be attributable to AMR, and as many as 8.22 million deaths could be associated with AMR annually under a reference scenario where resistance continues to rise without significant intervention (GBD 2021 Antimicrobial Resistance Collaborators, 2024). Conventional antibiotic therapy is becoming more ineffective due to resistance development, off-target effects, microbiome alteration, and toxicity concerns. These constraints demand novel molecular strategies capable of selectively recognising and neutralising harmful microorganisms while preventing the progression of resistance (Gupta et al., 2016; Jain et al., 2024; Jain et al., 2026). Nucleic acid-based therapeutics and diagnostics have drawn substantial interest as alternative molecular recognition agents, specifically aptamers (Ye et al., 2024).

Aptamers are short, single-stranded oligonucleotides (20-100 nucleotides) that fold into distinct three-dimensional structures, allowing them to bind preferentially to a variety of molecular targets. These targets include proteins, small molecules, whole cells, and even complex structures such as biofilms. These targets include proteins, small molecules, intact cells, and even complex structures such as biofilms. Aptamers are produced via the SELEX procedure, an iterative in vitro selection approach that enriches high-affinity binding sequences from vast random libraries. Aptamers offer several advantages over antibodies, including chemical synthesis without batch-to-batch variation, greater stability, ease of modification, reduced immunogenicity, and cost-effective synthesis.

### Antibacterial Mechanisms of Aptamers

The rapid emergence of multidrug-resistant (MDR) bacterial strains, driven by genetic mutations and indiscriminate antimicrobial use, has significantly reduced the efficacy of major antibiotic classes against pathogens such as Staphylococcus aureus, Pseudomonas aeruginosa, Escherichia coli, Salmonella typhi, and Mycobacterium tuberculosis. In response, antibacterial aptamers have emerged to target a broad spectrum of bacterial components and pathogenic processes, expanding their applicability from simple recognition molecules to multifunctional agents capable of therapeutic intervention and improved diagnostics (Figure 1).

**1. Therapeutic Mechanisms:** Aptamers exert their therapeutic effects either by directly inhibiting or suppressing mechanisms for bacterial survival and virulence or indirectly by delivering therapeutic drugs, molecules, antimicrobial peptides, siRNA, photherapeutic agents, etc, for enhancing the effects.

1. <u>Inhibiting Bacterial reproduction and survival:</u> Aptamers target critical proteins such as BamA in Pseudomonas aeruginosa, disrupting outer membrane protein assembly and compromising membrane integrity, leading to impaired colony formation and direct antimicrobial effects (Selvam et al., 2023). Similarly, aptamers targeting key metabolic or persistence-related enzymes, such as polyphosphate kinase 2 (PPK2) in Mycobacterium tuberculosis, that interfere with polyphosphate-dependent stress response and virulence pathways, thereby suppressing bacterial survival and replication (Shum et al., 2011).
2. <u>Suppress Biofilm Formation:</u> Aptamers bind and block adhesion factors (e.g., PBP2a), targeting the polysaccharide intercellular adhesin (PIA), disrupting quorum-sensing pathways, and altering extracellular matrix integrity, thereby inhibiting biofilm development and enhancing antibiotic susceptibility (Oroh et al., 2020; Matchawong et al., 2022).
3. <u>Neutralization of Bacterial Toxins (or Virulence factors):</u> Aptamers neuralize bacterial toxins such as α-toxin, staphylococcal enterotoxins (SEB, SEA), anthrax lethal factor (LF), and botulinum neurotoxin light chains, thereby preventing cytotoxicity, pore formation, inflammatory cytokine activation, and host cell damage (Vivekananda et al., 2014; Wang et al., 2015; Oliveira et al., 2024).
4. <u>Targeted Delivery via Nanoconjugates:</u> Aptamer-functionalized nanoparticles, including gold, silver, graphene oxide, liposomes and mesoporous silica, enable selective delivery of antibiotics and antimicrobial peptides to infected cells, improving intracellular targeting and reducing systemic toxicity. Additionally, aptamer-mediated delivery of siRNA/miRNA to regulate gene expression involved in virulence and resistance pathways. Similarly, in photothermal and photodynamic approaches, aptamer-conjugated nanomaterials generate localized heat or reactive oxygen species under light irradiation, causing oxidative damage to bacterial DNA, proteins, and membranes. Recently, radiolabeled targeted therapy involving aptamer conjugation to radionuclides (e.g., LJLJCu, ¹LJLJLu, ²²LJAc) for targeted radiation delivery represents a potential future antibacterial theranostic approach (Ye et al., 2024).
5. <u>Prevention of Bacterial Invasion into Host Immune Cells:</u> Intracellular pathogens evade immune clearance by entering the host immune cells. Aptamers targeting virulence-associated surface molecules, such as ManLAM in M. tuberculosis and type IVB pili in Salmonella typhimurium, inhibit bacterial entry into macrophages and monocytes, thereby enhancing host defense mechanisms (Chen et al., 2012).
6. <u>Modulation of Host Immune Response:</u> Beyond direct antibacterial activity, aptamers function as immunotherapeutic agents by modulating host immune pathways. They regulate Toll-like receptor signalling (e.g., TLR2 antagonism), enhance adaptive cytokine responses (e.g., IFN-γ, IL-15, IL-17), promote M1 macrophage polarization, and block immune checkpoints (e.g., PD-1), thereby restoring effective immune-mediated bacterial clearance (Afrasiabi et al., 2020).
**2.** Diagnostic application:

**Figure 1:**
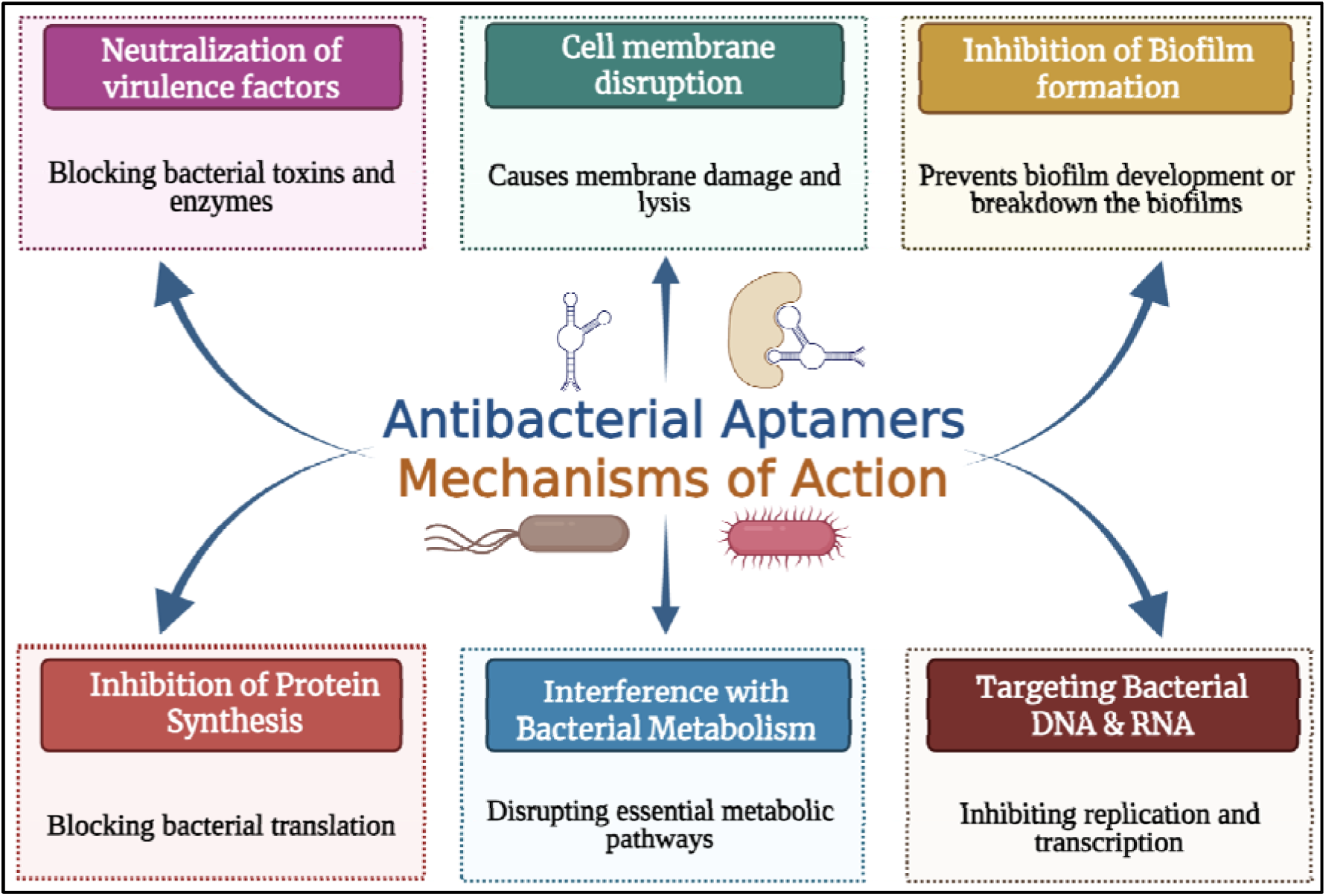
**General mechanism of action of antibacterial aptamers**

In parallel with therapeutic advancements, antibacterial aptamers have become powerful tools for detecting pathogens quickly and accurately. Conventional diagnostic methods, such as smear microscopy, culture-based assays, PCR, and immunoassays, have limited sensitivity, long processing times, or the inability to distinguish disease states, making them unsuitable for point-of-care (POC) applications, especially in complex samples like biofilms and aqueous environments.

Aptamers that have been produced against whole intact cells, surface proteins, and secreted antigens, enabling highly specific molecular recognition. These aptamers have been integrating into cutting-edge biosensing platforms, comprising nanoparticle-based electrochemical quartz crystal microbalance, fluorescence assays, surface-enhanced Raman spectroscopy, and amplification-based detection methods. Such systems produced enhanced detection limits, quick turnaround times, and a diagnosis time of roughly 2 hours while maintaining good specificity and sensitivity in clinical samples (Moulick and Bhattacharjee, 2020) (Chen et al., 2021; Chen et al., 2022; Ye et al., 2024).

### Need for centralized antibacterial aptamer repository

The success of antibacterial peptide research has been largely driven by the availability of specialized computational resources that systematically organize experimental data and assist predictive model development. Community databases, including recent versions of APD, CAMP, DRAMP, and DBAASP, have played critical roles in curating experimentally validated antimicrobial peptides, allowing for comparative analyses, computational modelling, and rational therapeutic design (Pirtskhalava et al., 2021; Gawde et al., 2023; Ma et al., 2025; Wang et al., 2026). More recently, comprehensive peptide databases like Peptipedia v2.0 (Cabas-Mora et al., 2024) and AI-driven computational frameworks have accelerated peptide identification, functional annotation, and activity prediction (Liu et al., 2026). Similarly, domain-specific resources, such as curated databases of experimentally validated anti-tubercular peptides, anti-tubercular peptide prediction tools, and open-source drug discovery platforms, have significantly advanced computational research against Mycobacterium tuberculosis and other bacterial pathogens (Bhardwaj et al., 2011; Usmani et al., 2018b; Usmani et al., 2018a; Bajiya et al., 2024; Roque-Borda et al., 2025; Wang et al., 2025). These examples clearly show how specialized databases and computational platforms help to standardize data, benchmark it, construct predictive models, and conduct translational antimicrobial research.

In contrast, computational resources dedicated to antibacterial aptamers remain limited. Several general-purpose aptamer databases, including the Aptamer Database (Lee et al., 2004), which was introduced as a collaborative knowledge base describing aptamer sequences and SELEX experiments and providing standardized experimental metadata.

Commercial and semi-public repositories, such as the Aptagen Aptamer Database, and curated platforms, such as Aptamer Atlas, have also helped consolidate aptamer-related information. Similarly, Most databases aggregate aptamers targeting diverse molecules, including small compounds, biomarkers, proteins, and eukaryotic receptors, without specialized categorization for bacterial species or antibacterial functionality. High-throughput SELEX analysis platforms such as AptaSUITE (Hoinka et al., 2018) and FASTAptamer (Kramer et al., 2022) have provided integrated pipelines for processing next-generation sequencing data from selection experiments. More recently, machine learning-, deep learning-, and large language-based approaches, such as DeepAptamer (Yang et al., 2025b), AptaTrans (Shin et al., 2023), and RaptScore (Kimura-Yamazaki et al., 2026), have been proposed to predict aptamer functionality from sequence features. While these resources are valuable, they are not specifically tailored to antibacterial applications. Furthermore, many existing repositories are not optimized for computational benchmarking and lack uniformly formatted datasets that facilitate machine learning model training or large-scale comparative analysis.

Despite the growing amount of aptamer research, the development of prediction models for antibacterial aptamers is constrained by the lack of a dedicated, standardised benchmark dataset. Experimentally verified antibacterial aptamers are scattered across individual articles and additional resources and are frequently described in diverse forms, with uneven annotation of sequence characteristics, target definitions, and binding affinities. This fragmentation impedes systematic data capture, adds bias into computational pipelines, and prevents meaningful benchmarking of machine learning and bioinformatics applications. Furthermore, despite promising preclinical studies demonstrating the therapeutic and diagnostic potential of antibacterial aptamers, no clinical trials involving antibacterial or bacteria-associated aptamers are currently registered on <u>ClinicalTrials.gov</u>, underscoring the gap between experimental research and clinical application.

To address these challenges, a centralized resource dedicated solely to bacterial targets can offer structured annotation of organism-specific information, target classification (whole cell, membrane protein, toxin, enzyme, or biofilm component), sequence characteristics, chemical modifications, and validated affinity parameters. Such standardization enables repeatability, large-scale benchmarking, comparative research, rational aptamer design, and data-driven predictive modeling. In this regard, we developed AptBacterialDB, a comprehensive, manually curated database that integrates experimentally demonstrated antibacterial aptamers into an organised, searchable, and computationally accessible framework. It aims to accelerate the systematic exploration of antibacterial aptamers and support the development of next-generation diagnostics and therapeutics.

## Materials and Methods

### 1. Collection of Data

Data for AptBacterialDB were obtained through a systematic literature review of peer-reviewed publications in major scientific repositories, including PubMed and Google Scholar. Multiple search combinations related to antibacterial aptamers or aptamers for bacteria were used to retrieve relevant studies. Only studies reporting experimentally validated antibacterial aptamers with clearly defined sequences and targets were included. Additionally, entries from existing databases, such as the UTexas Aptamer Database (Askari et al., 2024), AptaBase (https://www.iitg.ac.in/proj/aptabase/about.html), and AptaDB (Chen et al., 2024), were included. These search parameters led to a total of 470 PMIDs and 89 DOIs over the period from 1999 to 2026. All retrieved entries were manually curated to ensure accuracy and consistency. Redundant entries and duplicate records were carefully removed during the curation process. The entire database creation process, from data collection to development, is shown in Figure 2.

**Figure 2:**
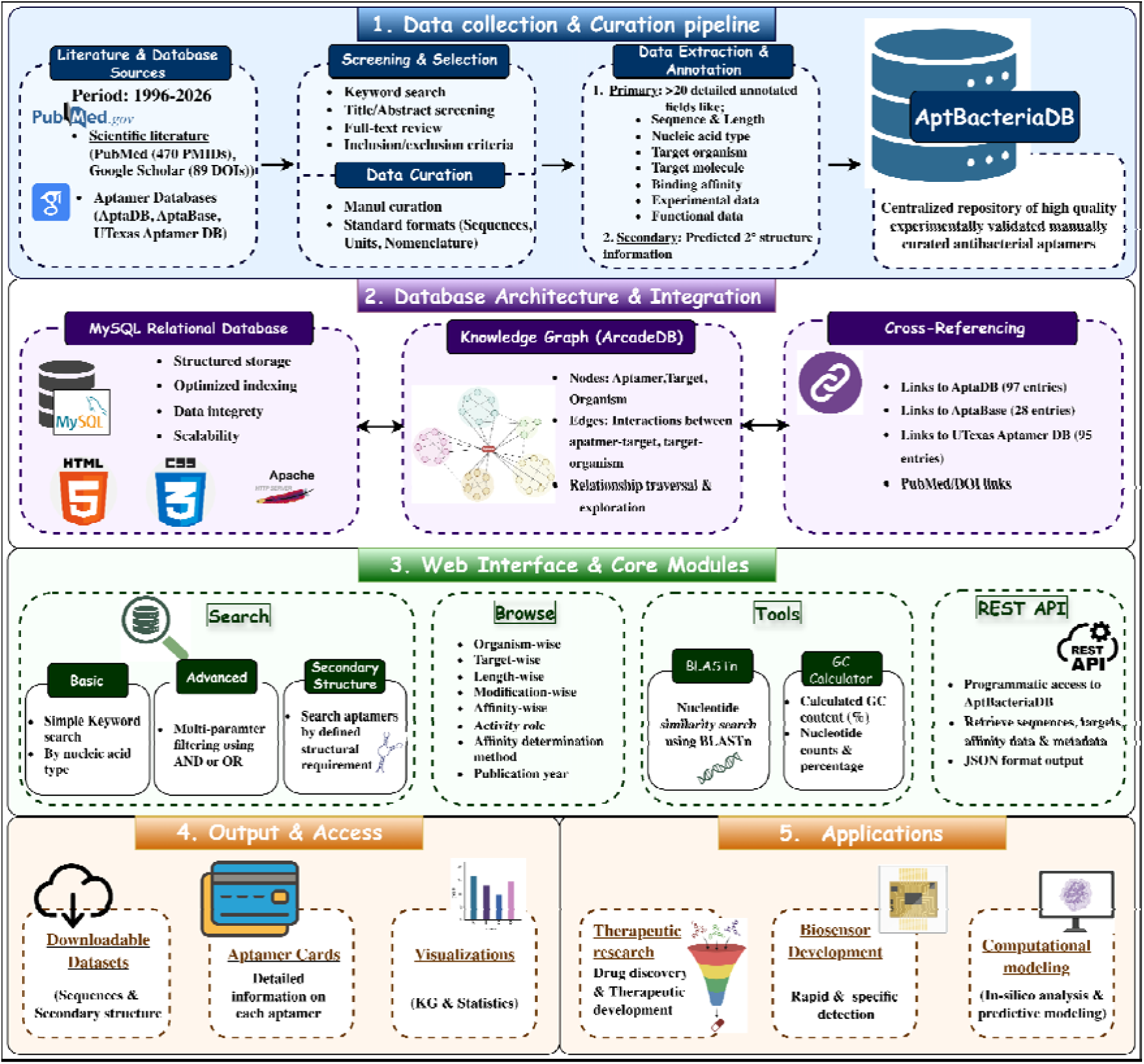
**The Architecture of AptBacterialDB**

### 2. Annotating the Data

Each aptamer entry was annotated with comprehensive metadata, including primary information on aptamer sequence and length, nucleic acid type (DNA or RNA), chemical modifications, target organism, aptamer target, SELEX methodology, reported binding affinity values, experimental validation techniques and activity role. Also, cytotoxicity, stability, half-life and patent information were reported where available. Secondary information, such as structural details, was obtained using the aptamer sequences. For that, we used RNAfold, a program in the ViennaRNA Package 2.0 (Lorenz et al., 2011), to predict aptamer secondary structures based on MFE values. RNAfold systematically evaluates possible base-pairing interactions to identify the thermodynamically most stable secondary structure, simultaneously reporting the predicted MFE values and providing dot-bracket notations for quick visualization. Also obtained the structural elements, including stems, loops, hairpins, and related structures. The entries have been linked to their corresponding aptamer cards, which contain this detailed, curated information.

### 3. Architecture of Database

The database was created using the MySQL (version 5.5.62) and Apache HTTP Server (version 2.4.7) frameworks to enable organised data storage and efficient retrieval. The backend was written in PHP, while the frontend was built with HTML5, CSS3, and JavaScript to deliver an interactive, responsive user experience. The database is mobile, tablet, and desktop friendly, with interconnected tables including aptamer characteristics, bacterial type, target information and experimental methods for querying, browsing, and analyzing data. Users can access it for free at https://webs.iiitd.edu.in/raghava/aptbacterialdb/.

### 4. Web Interface

AptBacterialDB web server, which is openly available, has been built to help the community. It provides information on antibacterial aptamers, their targets, and other relevant topics on a single platform. Multiple significant modules in the web server are provided for an effortless search facility, which are as follows;

**1. Search:** The Search module consists of three submodules: “Simple Search”, which allows users to perform quick queries using basic keywords such as bacterial organism name, aptamer ID, and target molecule. In this basic search module, the output can be adjusted based on the type of nucleic acid. The “Advanced Search” module enables multi-parameter filtering to support refined data retrieval. “Secondary Structure Search” allows the user to search aptamers based on the selected defined structural requirements.
**2. Browse:** The Browse module allows users to systematically explore the database by predefined categories. This module is particularly useful for users interested in investigating all aptamers associated with the specific browsing options “Selex method-wise’, ‘Target organism-wise’, ‘Target-wise’, ‘Length-wise’, Modification-wise’, ‘Activity role’, ‘Affinity Determination method’, and ‘Publication year’.
**3. Tools:** To enable similarity-based search, the nucleotide module of Basic Local Alignment Search Tool (BLASTn) has been implemented (Altschul et al., 1990). The user submits the FASTA format nucleotide sequence with default or specified settings to the BLAST module, and the server runs the BLAST search against the data stored in the database. Similarly, the GC calculator allows the user to obtain counts and percentages of all nucleotides, as well as the GC percentage.
**4. Knowledge Graph:** A Knowledge Graph (KG) module has been created using ArcadeDB [https://arcadedb.com/knowledge-graphs.html] to enable semantic organizing and relationship-based exploration of antibacterial aptamer data and openCypher, a declarative property graph query language. This module facilitates effective relationship-based querying and integrative analysis of antimicrobial aptamer interactions by allowing users to visualize and explore relationships between aptamers and bacterial organisms. The module also supports dynamic neighborhood expansion, allowing exploration of both direct and indirect associations among aptamers, targets, organisms, and functional applications.
**5. Cross-Referencing Module:** Because different repositories contain different types of information, we have linked entries that are also present in other repositories, including the UTexas Aptamer Database, AptaBase, and AptaDB, enabling users to access interdisciplinary information.
**6. Rest API:** To enable programmatic access to the database, we have provided a RESTful API. Users can retrieve aptamer sequences, target details, binding affinity data, and associated metadata in a structured JSON format. This feature facilitates automated data extraction and integration into computational pipelines for large-scale analysis and predictive modeling.

## Results and Discussion

AptBacterialDB comprises 2131 entries and is a comprehensive collection of experimentally verified antibacterial aptamers targeting a broad spectrum of Gram-positive and Gram-negative bacterial pathogens. The curated entries show increased interest in aptamer-based antibacterial methods and the variety of bacterial targets studied in recent years. Of which, 97 entries are also found in AptaDB, 29 in Aptabase, and 95 in the UTexas aptamer database. This repository includes 75 bacterial types and 124 reported aptamer targets, along with information on antibacterial aptamers up to 2026. The database contains 1555 unique aptamer sequences, 189 unique modifications, 40 different selection strategies, and 44 different affinity methods. Furthermore, the KG, which is graph-based and has nodes and edges, integrates connected data on aptamers, bacterial targets, and target species. Currently, the KG has 1,842 nodes and 4,853 edges, which represent biological entities and their empirically confirmed relationships. The cumulative growth analysis reveals a significant increase in antibacterial aptamer entries after 2014, with rapid expansion between 2014-2018 and 2019-2023, as shown in Figure 3, indicating increased global interest in alternative antimicrobial strategies in the face of rising antibiotic resistance.

**Figure 3:**
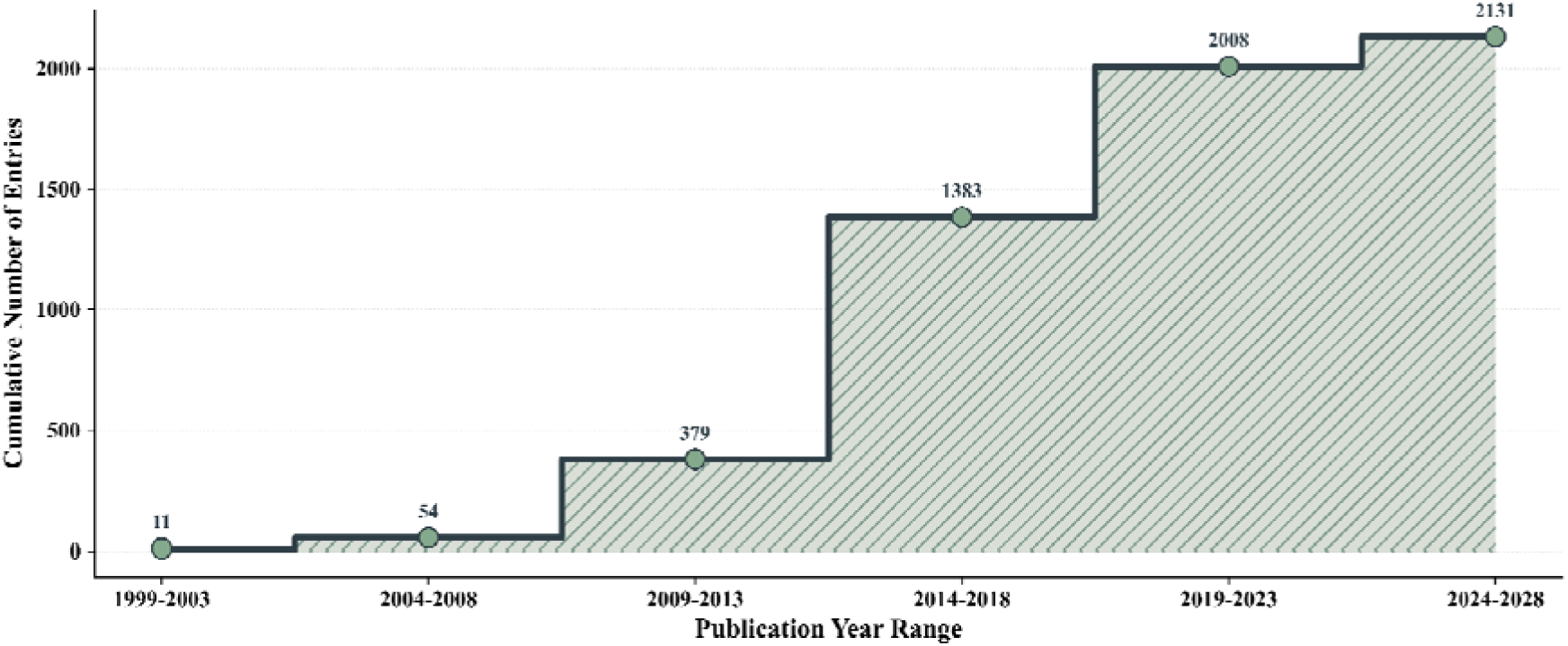
**Line plot showing the cumulative year-wise growth trend of aptamers in antibacterial studies**

The analysis of aptamer type in the antibacterial database reveals that ssDNA aptamers are dominant (96.1%). In contrast, ssRNA aptamers represent only 3.9% of the entries, since they are more chemically stable, cheaper to synthesise and easier to handle than RNA, which is more susceptible to nuclease degradation and often requires chemical modifications. Also, the entries based on the target organism, as shown in Figure 4, indicate that Escherichia coli (30.4%), Staphylococcus aureus (25.2%), and Mycobacterium tuberculosis (20.1%) together represent the majority of antibacterial aptamer targets. At the target level, whole-cell with an unspecific target (62.4 %) is the most common, followed by lipopolysaccharide (3.6 %) and outer membrane protein (1.9 %). In terms of length, aptamers are typically in the 61-80 nt length range, especially in biosensing and detection applications. Therapeutic roles are comparatively fewer and more evenly distributed across moderate length ranges.

**Figure 4:**
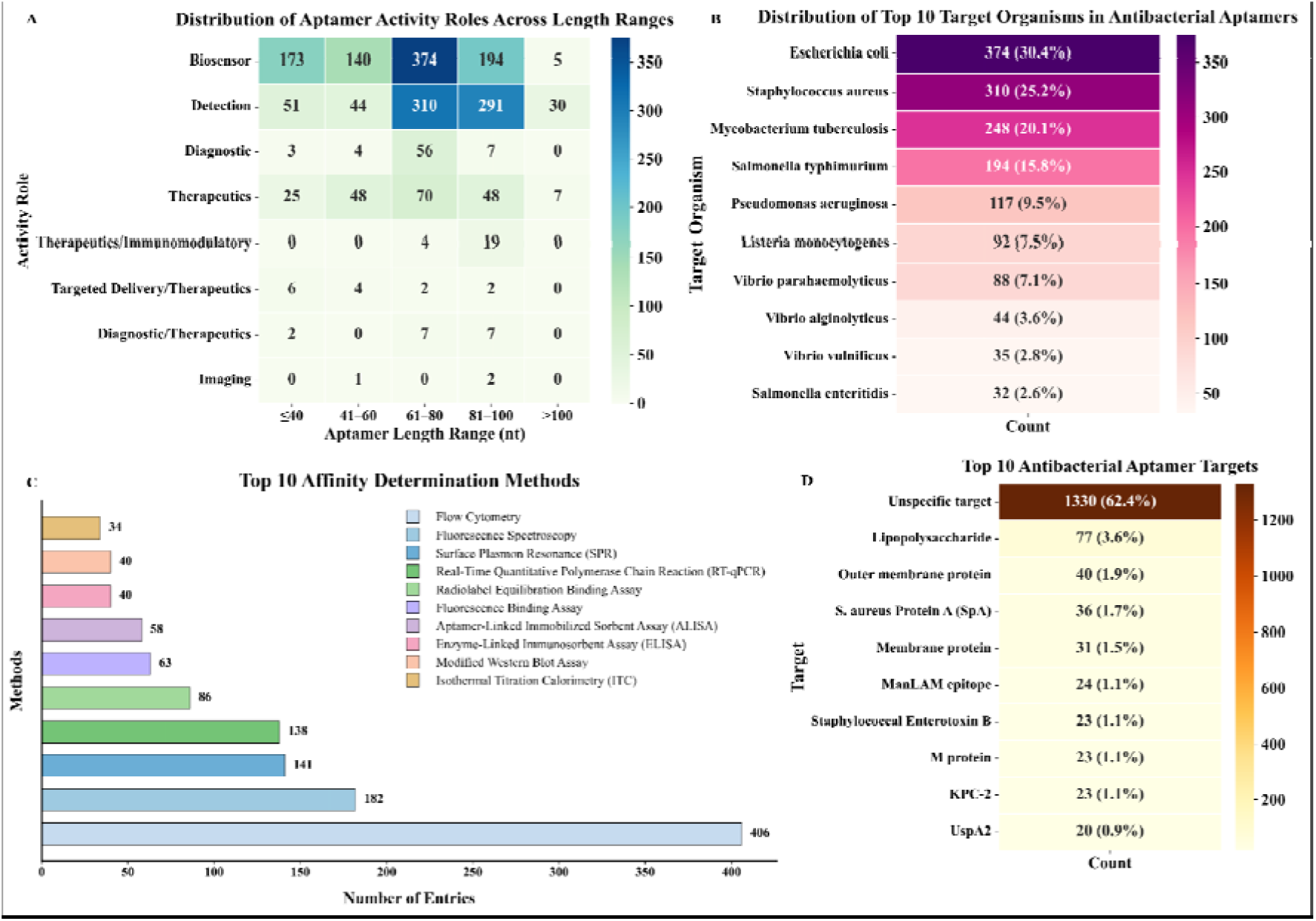
**Statistics of (A) Activity roles depending on aptamer length, (B) Top 10 bacterial target organisms, (C) Top 10 affinity determination techniques, (D) Top 10 bacterial targets for aptamer research in AptBacterialDB based on the number of entries.**

On analyzing the secondary structural elements, the stem regions have the highest percentage (∼44%), followed by multi-loops (∼30%) and internal loops (∼25%). This indicates that aptamers tend to have complex, stable conformations, with structured stem regions providing stability and loop regions aiding target binding. Moreover, the MFE distribution reveals that most aptamers have intermediate thermodynamic stability, with a subgroup exhibiting very stable folded structures, emphasising the structural variability in the database. The majority of the aptamers displayed high stability, with MFE values ranging from −5 to −12 kcal/mol and a negatively skewed distribution (Figure 5). A fraction of aptamers showed significant negative values for MFE values (<-20 kcal/mol), suggesting the presence of tightly folded structures. Overall, the observed distribution indicates that aptamers maintain a balance between structural stability and flexibility, both of which are important for effective target binding and functional activity (Yang et al., 2025a).

**Figure 5:**
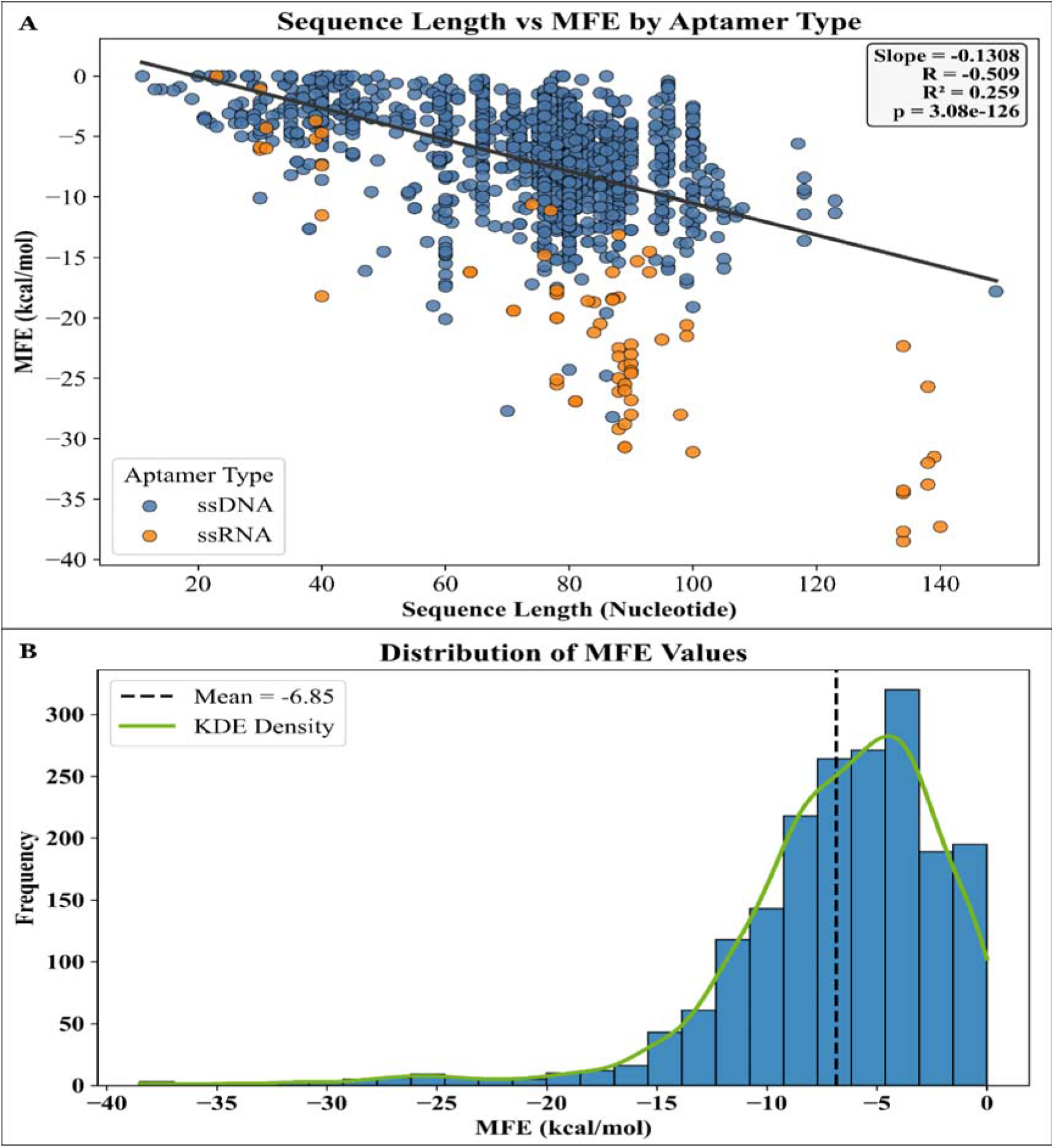
**(A) A regression plot of projected MFE values against aptamer sequence length. (B) Bar plot depicting predicted MFE values throughout the antibacterial aptamer dataset.**

## Conclusion

AptBacterialDB (https://webs.iiitd.edu.in/raghava/aptbacterialdb) is a large, manually curated collection of experimentally characterised antibacterial aptamers, including 2131 aptamers targeting approx 75 different bacterial classes, and 124 aptamer targets. By combining sequence-level data with extensive experimental annotations and application classifications, the database addresses a major gap in antibacterial research. It offers a comprehensive and user-friendly platform for researchers working on molecular therapeutics, biosensor development, antimicrobial drug delivery, and computational modeling. This will aid in the development of prediction algorithms for identifying antibacterial aptamers.

## Data Availability

We will update it soon.

## Authors’ contributions

NB manually collected, curated and analysed the data. IG developed the backend and frontend of the web server. NB and GPSR prepared the manuscript. The manuscript was reviewed by NB, IG and GPSR. The project was conceived and coordinated by GPSR. The final manuscript was read and approved by all the authors.

## Declaration of generative AI and AI-assisted technologies

During the preparation of this work, the author(s) used ChatGPT in order to improve the writing and language of the manuscript. After using this tool/service, the author(s) reviewed and edited the content as needed and take(s) full responsibility for the content of the published article.

